# Ozone nanobubble treatment effectively reduced pathogenic Gram positive and negative bacteria in freshwater and safe for tilapia

**DOI:** 10.1101/2020.06.07.138297

**Authors:** Chayuda Jhunkeaw, Nareerat Khongcharoen, Naruporn Rungrueng, Pattiya Sangpo, Wattana Panphut, Anat Thapinta, Saengchan Senapin, Sophie St-Hilaire, Ha Thanh Dong

## Abstract

High concentrations of pathogenic bacteria in water usually results in outbreaks of bacterial diseases in farmed fish. Here, we explored the potential application of an emerging nanobubble technology in freshwater aquaculture. Specifically, we aimed to determine if this technology was effective at reducing the concentration of pathogenic bacteria in the water, and to assess whether it was safe for fish. An ozone nanobubble (NB-O_3_) treatment protocol was established based on examination of nanobubble size, concentration, disinfection property, and impact on fish health. A 10-min treatment with NB-O_3_ in 50 L water generated approximately 2-3 × 10^7^ bubbles with majority sizes less than 130 nm and ozone level of ∼800 mV ORP. A single treatment with water contaminated with either *Streptococcus agalactiae* or *Aeromonas veronii* effectively reduced 96.11-97.92 % of the bacterial load. This same protocol was repeated 3 times with 99.93-99.99 % reduction in the bacterial concentration. In comparison, bacterial concentration the control tanks remained the same level during the experiments. In fish-cultured water with the presence of organic matter (e.g. mucus, feces, bacterial flora, feed, etc.), the disinfection property of NB-O_3_ was reduced i.e bacterial concentration was reduced by 42.94 %, 84.94 % and 99.27 % after the first, second and third treatments, respectively. To evaluate the safety of NB-O_3_ to fish, juvenile Nile tilapia were exposed to NB-O_3_ treatment for 10 minutes. No mortality was observed during the treatment or 48 h post treatment. Gill histology examination revealed that a single NB-O_3_ treatment caused no alteration morphology. However, damage in the gill filaments was noticed in the fish receiving two or three consecutive exposures within the same day. Results of all the experiments conducted in this study suggest that NB-O_3_ technology is promising for controlling pathogenic bacteria in aquaculture systems, and may be useful at reducing the risk of bacterial disease outbreaks in farmed fish.

## Introduction

The aquaculture sector has played a vital role in global food security. It supplies protein for approximately 4.5 billion peoples and employs 19.3 million people worldwide (Béné et al., 2015; FAO, 2018). Similar to other food sectors, aquaculture has faced increasing challenges with infectious diseases. Control of these diseases has led to an increase in the use of antimicrobials (Watts et al., 2017; World Bank, 2014). Of particular importance to public health has been the increase in antimicrobial resistance (AMR). Alternatives for these products to control bacterial infections in all food production sectors have increased over the last few years (Reverter et al., 2020; Watts et al., 2017). In the aquaculture sector, previous and current approaches focus mainly on antibacterial compounds derived from natural products, probiotics, immunostimulants, and vaccines for prevention strategies (Reverter et al., 2020; Watts et al., 2017).

Other prevention strategies, usually used in closed recirculating systems to reduce the bacterial concentration that fish are exposed to, include water treatments with UV or Ozone. Both of these treatments have issues for the aquaculture industry. UV requires that water be very clean when it is exposed to the light source, which renders it less than ideal in pond culture. Ozone has a low dissolution property, rapid decomposition in water and can be lethal to fish (Huyben et al., 2018; Xia et al., 2019). More effective non-chemical water treatment technology is needed to improve water quality for aquaculture systems such as intensive pond culture systems.

Nanobubble technology is an emerging technology for wastewater treatment (Agarwal et al., 2011; Yamasaki et al., 2005) and recently being applied in aquaculture for the increasing concentration of dissolved oxygen in intensive aquaculture systems (Agarwal et al., 2011; Anzai et al., 2019; Mahasri et al., 2018; Rahmawati et al., 2020). This technology involves the injection of nano or ultrafine bubbles with a chosen gas into water (Agarwal et al., 2011; Anzai et al., 2019). Unlike macro- and microbubbles, these nanobubbles with a diameter less than 200 nm, have neutral buoyancy, thus remain in water for days (Agarwal et al., 2011; Takahashi et al., 2007).

Kurita et al. (2017) investigated the effect of exposing parasitic planktonic crustaceans to nanobubbles created from ozone (NB-O_3_). They reported that a 25 min treatment with NB-O_3_ successfully reduced 63% of the parasites compared to the untreated group. Most importantly, this treatment condition was safe for both sea cucumbers (*Apostichopus japonicas*) and sea urchins (*Strongylocentrotus intermedius*), which are commonly infected with these pathogenic crustaceans in Japanese aquaculture systems. In another study, Imaizumi et al. (2018) reported that NB-O_3_ could be used for disinfection of *Vibrio parahaemolyticus*, a unique strain causing early mortality syndrome/acute hepatopancreatic necrosis disease (EMS/AHPND) in whiteleg shrimp (*Penaeus vannamei*). However, in their study NB-O_3_ showed a negative effect on shrimp when administered at a high level (970 mV ORP). When the NB-O_3_ treated water was diluted by 50% and the results revealed that all shrimp exposing to pathogenic *V. parahaemolyticus* survived from the bacterial infection, while all shrimp died in the group without the NB-O_3_ treatment (Imaizumi et al., 2018).

Preliminary results of NB-O_3_ in marine aquaculture is promising. The impact of nanobubbles in water of different salinity suggests that this technology may be even more effective in fresh water (Li et al., 2013). However, there is a lack of studies on its effect on fresh water fish and their pathogens. This study aims at the assess whether NB-O_3_ can be used on fresh water fish pathogens and is safe for tilapia.

## Materials and Methods

### Concentration and size of nanobubbles

Two trials were carried out separately using the nanobubble generator (model: aQua+075MO; maker: AquaPro Solutions Pte Ltd, Singapore) to determine the size of the air and oxygen nanobubbles. The generator was operated in 100 L-fiberglass tanks containing 50 L distilled water for 30 min, with either natural air or oxygen gas with a flow rate of 1 L/min. 50 mL of water was sampled from each tank at 10, 15, 20, and 30 min. Water samples prior to the addition of nanobubbles were used as baseline standards. The concentration and size of nanobubbles were determined by NanoSight NS300 (Malvern Panalytical Ltd) with three replicates for each sample. Ozone nanobubble measurement was not done due to its oxidation effect on the NanoSight machine.

### Effect of ozone nanobubbles (NB-O_3_) treatment on water parameters

The experiment was performed in two separate tanks to evaluate the effect of NB-O_3_ on water parameters. Each tank contained 50 L of de-chlorinated tap water. Nanobubble generator was operated for 10 min in each tank. The temperature in degree Celsius (T^°^), dissolved oxygen (DO), pH and oxidation reduction potential (ORP) were measured using a multi-parameter meter (YSI Professional Plus) every 1-2 min during 10 min-run and 15 min after stopping the nanobubble generator.

### Bacterial isolates and growth conditions

The Gram-positive bacterium *Streptococcus agalactiae* isolated from a tilapia farm which was experiencing an outbreak of Streptococcosis, and Gram negative bacterium *Aeromonas veronii* associated with hemorrhagic septicemia in tilapia (Dong et al., 2017) were used in this study. Prior to experiments, the bacterial isolates were recovered from bacterial stocks stored at −80 ^°^C using tryptic soy agar (TSA) medium, incubated at 30 ^°^C. To prepare bacterial inoculum, single bacterial colonies were inoculated in 10 mL of tryptic soy broth (TSB) overnight at 30 ^°^C on a shaker platform (150 rpm). Five mL of bacterial culture was then sub-cultured in 500 mL of TSB, incubated with gentle shaking (150 rpm) at 30 °C until OD_600_ reached 0.8 (equivalent to ∼10^8^ CFU/mL). For subsequent trials, 100 mL of the bacterial culture was added into a tank containing 50 L de-chlorinated tap water.

### Effect of treatment time on disinfection property of NB-O_3_

An initial trial was carried out to investigate the effect of treatment time on the disinfection property of NB-O_3_. *S. agalactiae* was used as a representative bacterium in this time-course trial. The experiment was performed in two 100 L fiberglass tanks containing 50 L of de-chlorinated tap water each mixed with 100 mL bacterial culture (OD_600_ = 0.8). One tank was treated with NB-O_3_ while another tank was served as a control without NB-O_3_. Water was sampled from the four corners and the center of the tank (1 mL per spot). The samples were pooled together for conventional plate count enumeration at different time points. Samples were collect prior to inoculation (0 min), during treatment (5, 10 and 15 min) and after treatment (5, 10, and 15 min). The samples were 10 fold-serially diluted with sterile saline solution (NaCl 0.85%) and 100 µL of each dilution was spread on TSA in duplicates, incubated at 30 ^°^C for 36 h. Dilutions with a number of colonies ranging from 30-300 were used for counting and mean bacterial colonies of two replicate plates were calculated and expressed as CFU/mL. The percentage of bacterial reduction was calculated based on the formula below.

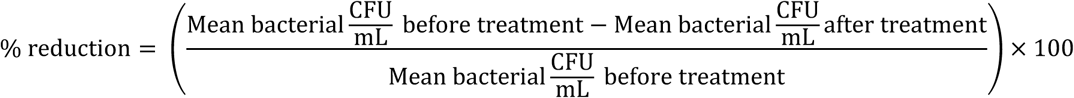

We compared the reduction in bacterial concentration in the tank exposed to ozone and the control tank for differences.

### Effect of NB-O_3_ on pathogenic Gram-positive and Gram-negative bacteria

To evaluate the effect of NB-O_3_ on bacterial pathogens of tilapia, *S. agalactiae* and *A. veronii* were used as representative Gram positive and Gram-negative bacteria, respectively. Each set of experiment comprised of 1 control tank (having normal aerator) and 3 treatment tanks (10 min-treating with NB-O_3_ 1 to 3 times at 15 min intervals). Each tank containing 50 L de-chlorinated tap water was mixed with 100 mL of bacterial suspension (OD_600_ = 0.8) as described above. Water was sampled from control and treatment tanks before (0 min) and 15 min after the end of each treatment to establish the bacterial concentration and the percentage of bacterial reduction. Temperature, pH, DO and ORP were also recorded during the experiment.

To investigate the ultrastructure of bacteria before and after treatment with NB-O_3_, two experimental tanks were set up in the same manner as the aforementioned treatment tanks, one tank contained *S. agalactiae* and the other contained *A. veronii*. Each tank was treated with NB-O_3_ for 10 min. Water (200 ml) was collected and concentrated to a 0.5 mL suspension before and 15 minutes after the NB-0_3_ treatment. The bacterial suspension was smeared on coverslips coated with Poly-L-lysine (Sigma-Aldrich) and air-dried for 3 hrs. The samples were subsequently fixed with glutaraldehyde 2.5% and 1% osmium tetroxide before dehydration with ethanol as described by Thanomsub et al. (2002). The ultrastructure of the bacteria was examined and photographed under a scanning electron microscope (SEM) (SU8000, Japan) operated at 10 kV.

### Effect of NB-O_3_ treatment on total bacteria in fish-culture water

Investigation of the disinfection property of NB-O_3_ was also evaluated using “culture” water (water from the fish-culture tanks which contained organic matter e.g. fish feces, mucus, left over feed and unknown aquatic bacterial flora). Fish-culture water was taken from tanks containing juvenile Nile tilapia (*Oreochromis niloticus*). A trial using three 10 min-NB-O3 exposure times administered 15 minutes apart was applied in to three fiberglass tanks containing 50 L fish-cultured water each. Water sampling scheme for total bacterial counts was conducted before and 15 min after the end of each treatment. Water temperature, pH, DO and ORP were monitored.

### Effect of NB-O_3_ on fish health and gill morphology

Animal use protocol in this study was granted by the Thai Institutional Animal Care and Use Committee (MUSC62-039-503). To investigate whether NB-O_3_ treatment had negative effects on gill morphology and fish life, we carried out a trial which included 2 control and 2 treatment tanks, each tank containing 20 apparently healthy *O. niloticus* juveniles of 6-8 g body weight. The 100 L fiberglass tanks had 50 L of de-chlorinated tap water. For the treatment tanks, NB-O_3_ was applied at 15 minute intervals 3 times for 10 minutes. The control tanks were treated with normal aeration. Two fish from each tank were randomly sampled after every treatment for wet-mount examination and histological study of the gills and the remaining fish were monitored for 48 h. For histological analysis, gill arches from one side of each fish were preserved in 10% neutral buffer formalin with a ratio of 1 sample/10 fixative (v/v) for 24 h before being placed in 70% ethanol for storage. The samples were then processed for routine histology and stained with hematoxylin and eosin (H&E). Gill morphology of the experimental fish was examined under a microscope equipped with a digital camera. We compared fish behavior, the gills of treated and untreated fish visually. Fish were also monitored for mortality over a period of 48 hours post treatment.

## Results

### Concentration and size of nanobubbles

The results of NanoSight readings from the air nanobubbles (NB-Air) (Fig. 1A) and the oxygen nanobubbles (NB-O_2_) (Fig. 1B) were similar. Majority of nanobubbles (or particles) were less than 130 nn in size. The concentration of these bubbles after a 10 min treatment was of 2.39 × 10^7^ ± 1.01 × 10^7^ particles/mL for NB-Air and 3.03 × 10^7^ ± 1.11 × 10^6^ particles/mL for NB-O_2_. Increasing treatment times (15, 20 and 30 min) generated larger bubbles with quantity in the same order of magnitude (Fig. 1). The result confirmed that the nanobubbler used in this study produced nanobubbles and 10 min operation in 50 L of water generated the most uniform nano-sizes. Thus, this scheme was also applied to generate ozone nanobubbles (NB-O_3_).

**Figure 1:**
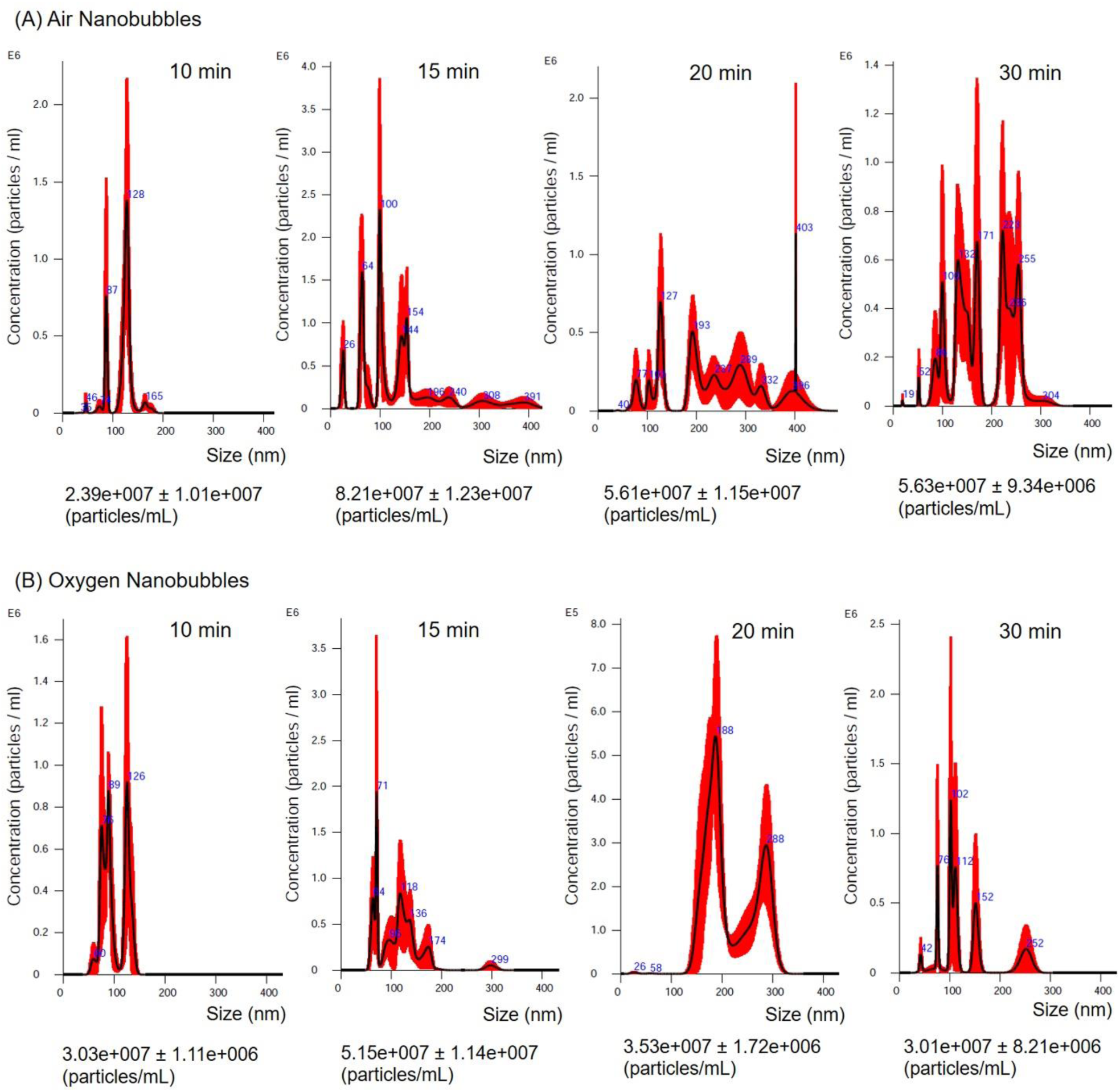
Concentration and size of bubbles generated using air (A) or oxygen (B) following treatment for 10, 15, 20 and 30 min. Peaks represent the concentration of dominant bubbles with similar sizes and blue numbers indicate the bubble sizes. Total concentrations of bubbles are shown at the bottom of each graph. Values were calculated from 3 replicate experiments.

### Effect of NB-O_3_ treatment on water parameters

Changes of water parameters (T^°^, DO, pH and ORP) during and after treatment with NB-O_3_ were consistently similar between the two trials (Fig. 2). Significant changes were observed in DO and ORP values while T° increased slightly (∼2°C) and pH remained relatively stable during and after NB-O_3_ treatment. With respect to DO, the value increased rapidly reaching to 23-25 mg/L after 10 min treatment and reduced slowly to ∼20 mg/L 15 min post treatment. By contrast, ORP increased quickly, reaching over 700 mV within 6 min and ∼800 mV within 10 min and dropped back to the starting level (∼300 mV) 15 min post treatment.

**Figure 2:**
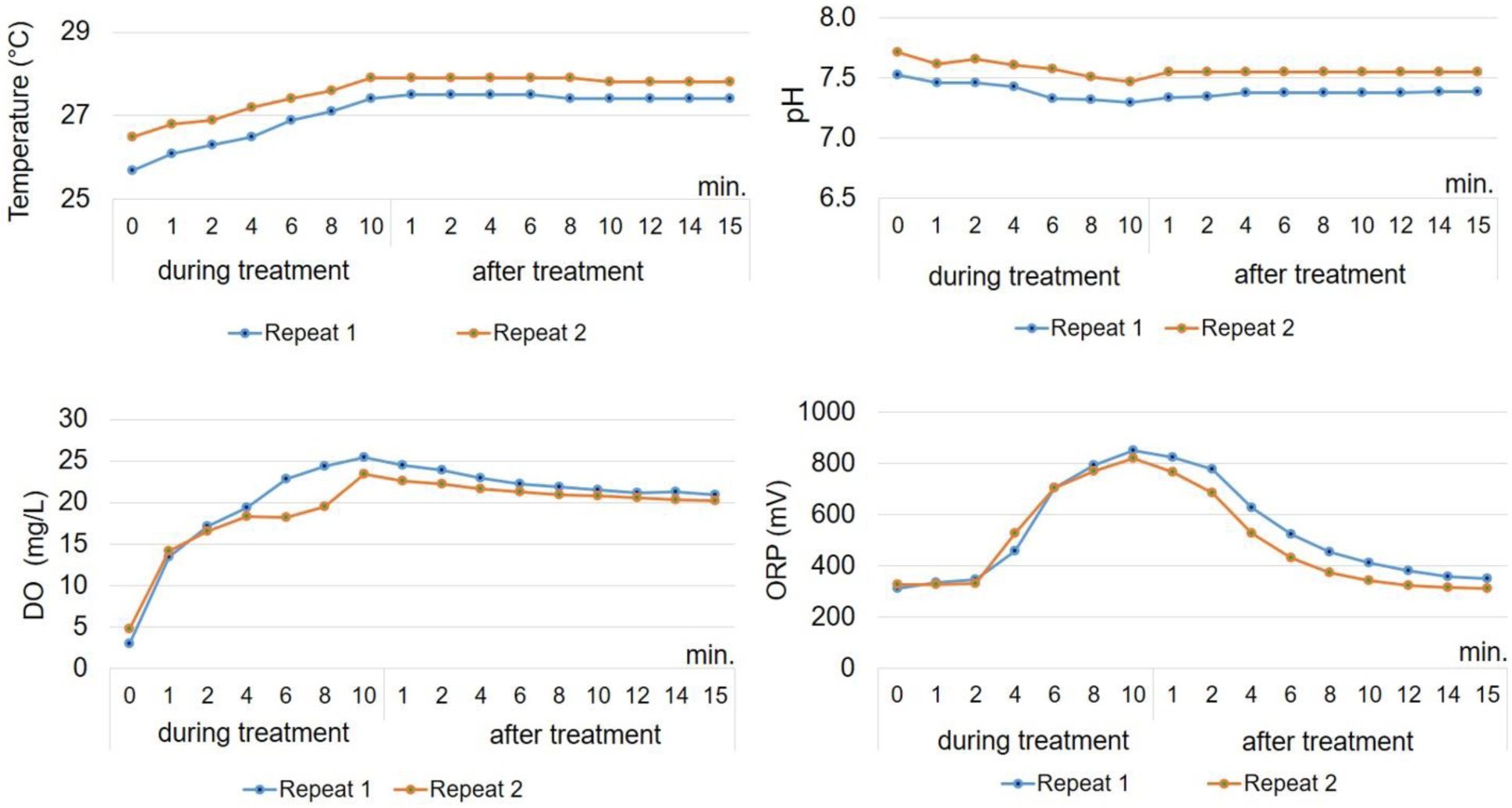
Water parameters (temperature, pH, DO and ORP) during 10 min treatment and 15 min after exposure to ozone nanobubbles. The experiment was carried out in 2 replicates.

### A 10-min NB-O_3_ treatment reduced >90% bacterial loads in water

As shown in Fig. 3, similar bacterial loads (*S. agalactiae*) at the starting point were used in the control tank (1.17 × 10^6^/mL) and treatment tank (1.83 × 10^6^/mL). However, upon NB-O_3_ treatment, bacterial density reduced quickly during exposure time in the treatment tank. The percentage drop in concentration in the treated group during the treatment at 5, 10 and 15 min were 62.30%, 97.76% and 99.40%, respectively, indicating that disinfection occurred rapidly during the treatment process. Bacterial concentration remained low 15 min after treatment. In contrast, bacterial concentration in the control tank remained stable at ∼10^6^ CFU/mL during the same time period (Fig. 3). With respect to water quality, changes were observed only in the treatment tank. DO increased from 6.2 mg/L (before treatment) to 21.8 mg/L (at 5 min), 25.8 mg/L (at 10 min) and 27.9 mg/L (at 15 min) and dropped to 23.3 mg/L at 15 min post treatment. Water temperature increased approximately 1 ^°^C every 5 min of the treatment, from 26.5 ^°^C (before treatment) to 29.2 ^°^C (at 15 min) and remained at this temperature 15 min post treatment. Relatively no change was observed in pH (7.6-7.7) and ORP (293-306 mV) during the experiment.

**Figure 3:**
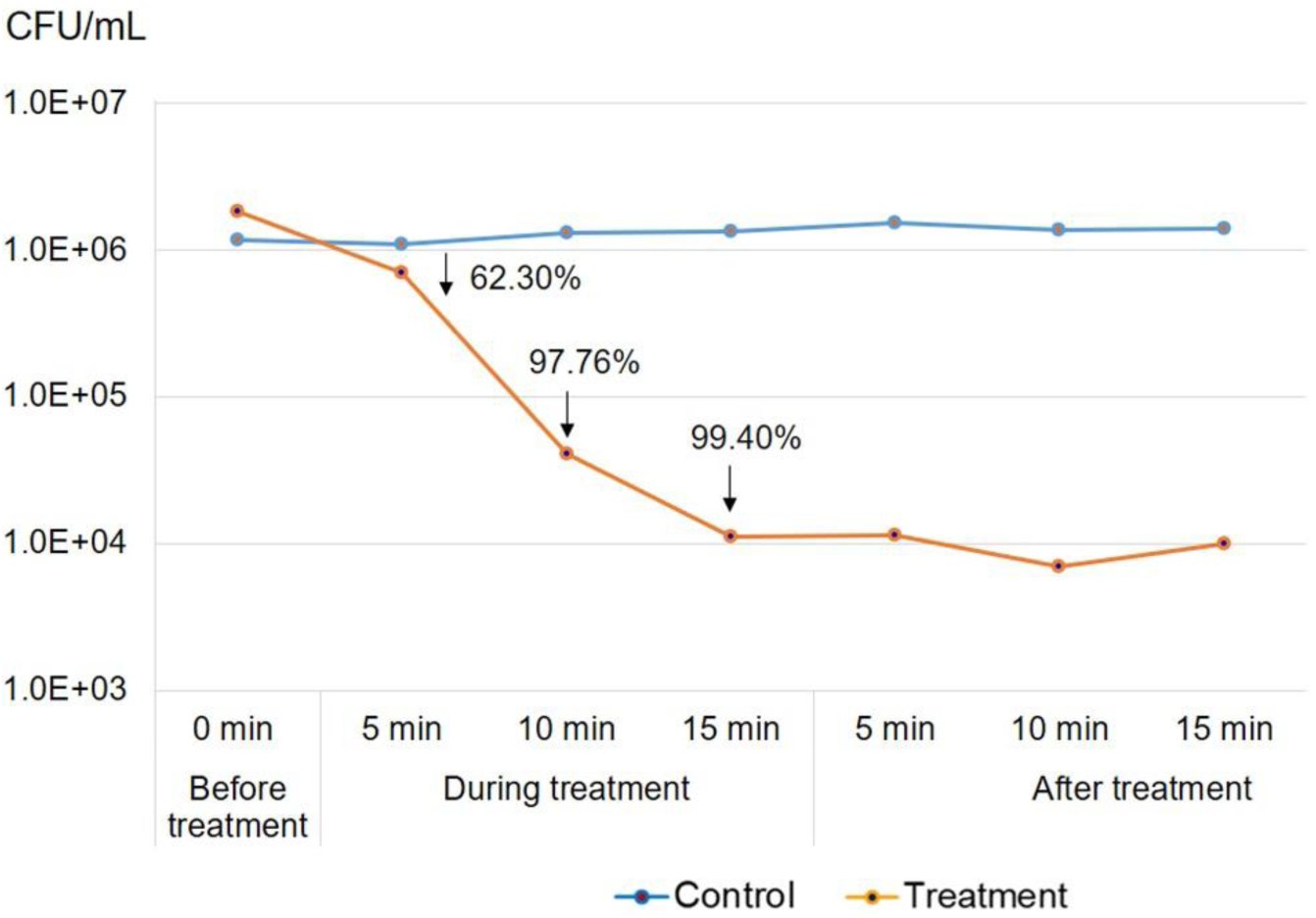
Number of *S. agalactiae* colony counts from the water with and without NB-O_3_ exposure (treatment and control group, respectively). NB-O_3_ treatment was performed for 15 min and stopped for 15 min. The water sample was collected from both the control and treatment groups every 5 min for plate count. Arrows indicated significant % reduction of bacterial counts compared to the starting point of the NB-O_3_ treatment group

### NB-O_3_ treatment effectively reduced both pathogenic Gram positive and negative bacteria

The trial with *S. agalactiae* started with similar bacterial loads; 1.17 × 10^6^ CFU/mL in the control tank and 3.45×10^6^ CFU/mL in treatment tanks (Fig. 4A). A single 10-min treatment with NB-O_3_ effectively reduced 96.11% bacterial load in the tank. When the same protocol was repeated for the second and third time, 99.93% and 99.99% bacteria were inactivated, respectively. The bacterial concentration in the control tank (without the NB-O_3_ treatment) was maintained at 10^6^ CFU/mL (Fig. 4A). Similar patterns were also observed in the trials with the Gram negative bacterium *A. veronii*. Average initial bacterial counts of *A. veronii* for control and treatment tanks were 1.03 × 10^6^ CFU/mL and 1.65 × 10^6^ CFU/mL, respectively. Following the 1^st^, 2^nd^ and 3^rd^ NB-O_3_ exposure, bacterial loads were reduced by 97.92, 99.99 and 99.99%, respectively (Fig. 4B). No significant changes in bacterial counts were observed in the control tank during the experiment (Fig. 4B). Changes in water quality were shown in Table 1. Temperature changes in the NB-O_3_ treatment tanks were 1.9-2.6 °C after the 1^st^ treatment, and 4.3-4.7 °C after 3^rd^ treatment, whereas pH values were relatively stable at 7.4 to 8.0. Notably, DO increased sharply (from 3.9-4.4 to 26.4-29.9 mg/L) and was maintained at this high level after every treatment, while ORP values did not increase as much as seen in the water study without bacteria (Fig. 2).

**Table 1:** Comparative water parameters in control and NB-O_3_ treatment groups with the presence of either *S. agalactiae* or *A. veronii* in the water

| Parameter | Measurement time | *S. agalactiae* |  | *A. veronii* |  |
| --- | --- | --- | --- | --- | --- |
|  |  | Control | NB-O<sub>3</sub> treatment | Control | NB-O<sub>3</sub> treatment |
| T<sup>0</sup> | Before treatment | 26.9 | 27.2 ± 0.3 | 27.5 | 25.9 ± 0.8 |
|  | 10 min (1<sup>st</sup>) | 26.9 | 29.8 ± 1.3 | 27.4 | 27.8 ± 0.6 |
|  | 10 min (2<sup>nd</sup>) | 27.0 | 30.4 ± 0.2 | 27.3 | 29.3 ± 0.6 |
|  | 10 min (3<sup>rd</sup>) | 27.0 | 31.5 ± 0.3 | 27.4 | 30.6 ± 0.5 |
| DO | Before treatment | 4.3 | 3.9 ± 0.5 | 4.7 | 4.4 ± 0.2 |
|  | 10 min (1<sup>st</sup>) | 4.3 | 27.8 ± 1.6 | 4.6 | 30.3 ± 2.4 |
|  | 10 min (2<sup>nd</sup>) | 4.2 | 26.9 ± 0.4 | 4.6 | 29.9 ± 0.1 |
|  | 10 min (3<sup>rd</sup>) | 4.2 | 26.4 ± 0.6 | 4.5 | 29.5 ± 1.0 |
| pH | Before treatment | 7.8 | 7.6 ± 0.2 | 7.8 | 8.0 ± 0.1 |
|  | 10 min (1<sup>st</sup>) | 7.8 | 7.5 ± 0.0 | 8.0 | 7.8 ± 0.1 |
|  | 10 min (2<sup>nd</sup>) | 7.8 | 7.4 ± 0.0 | 7.9 | 7.7 ± 0.1 |
|  | 10 min (3<sup>rd</sup>) | 7.8 | 7.4 ± 0.0 | 7.9 | 7.6 ± 0.0 |
| ORP | Before treatment | 325 | 290 ± 16 | 279 | 294 ± 6 |
|  | 10 min (1<sup>st</sup>) | 314 | 281 ± 7 | 289 | 271 ± 8 |
|  | 10 min (2<sup>nd</sup>) | 306 | 275 ± 4 | 261 | 270 ± 6 |
|  | 10 min (3<sup>rd</sup>) | 304 | 273 ± 3 | 265 | 272 ± 4 |

T<sup>0</sup>, temperature in degree Celsius; DO, dissolved oxygen; ORP, oxidation reduction potential.
Values in the NB-O<sub>3</sub> treatment are expressed as mean ± SD from 3 replicates.

**Figure 4:**
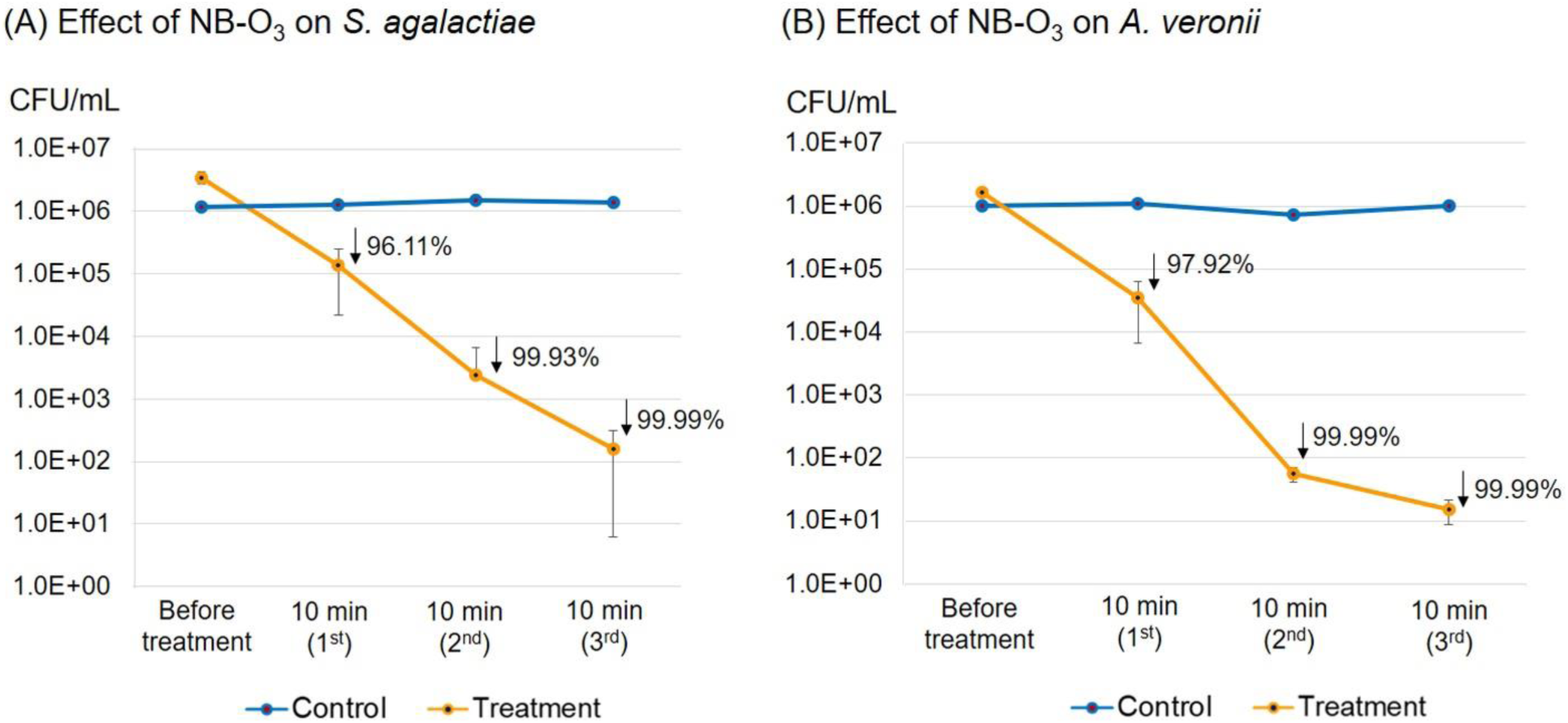
Bacterial counts of *S. agalactiae* (A) and *A. veronii* (B) upon exposure to NB-O_3_ 10 min three times continuously (orange lines) compared to that of the control water without NB-O_3_ (blue lines). Arrows indicated % reduction of bacterial loads compared to the starting bacterial concentration. Bars represent standard deviation from 3 replicates.

Ultrastructural examination of the bacterial surface by SEM revealed that the majority of bacterial cells (both *S. agalactiae* and *A. veronii*) were collapsed and destroyed after treatment with NB-O_3_ for 10 min compared to the normal intact surface structure of bacteria before treatment (Fig. 5).

**Figure 5:**
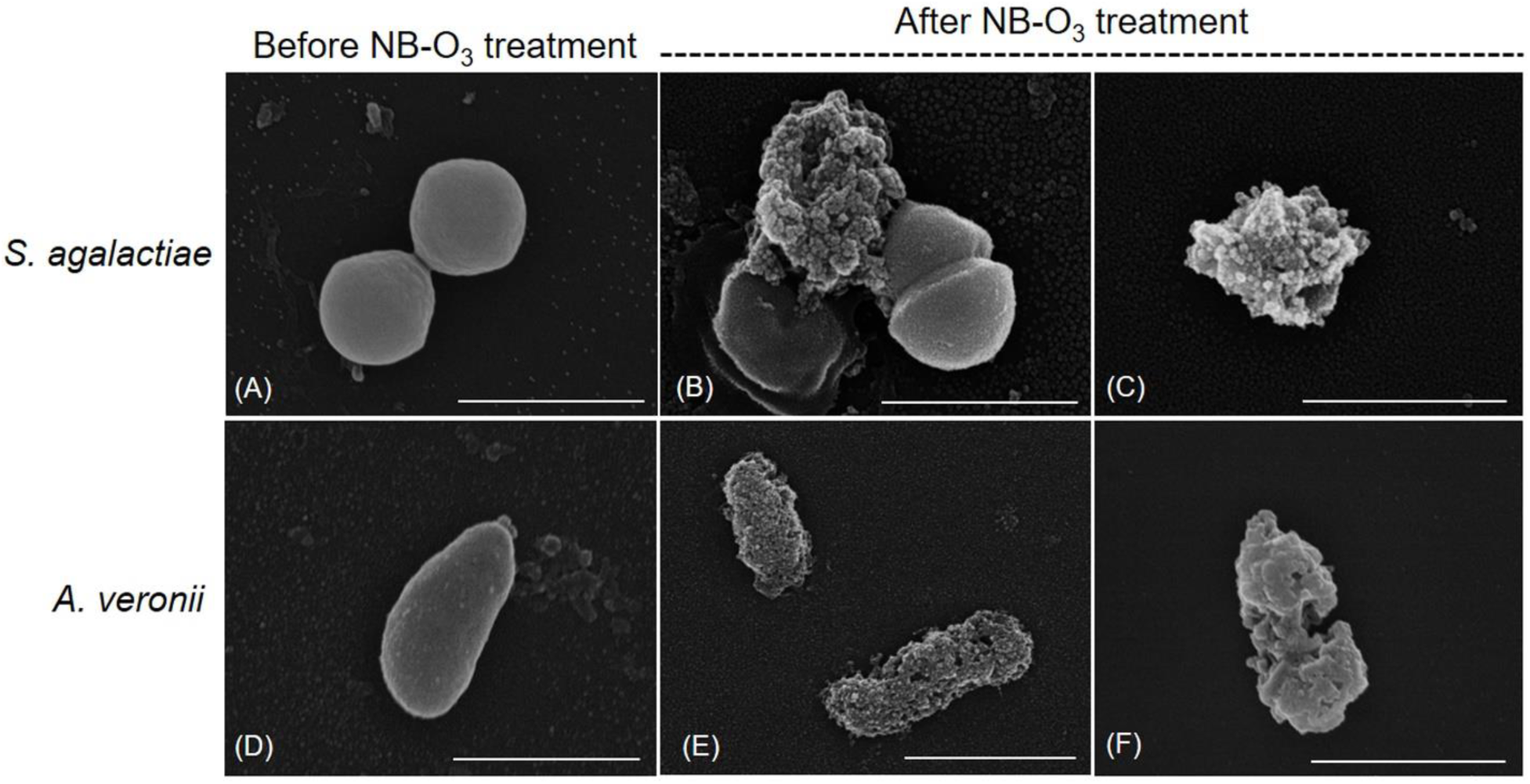
Scanning electron micrographs of *S. agalactiae* (A-C) and *A. veronii* (D-F) before and after treatment with NB-O_3_ for 10 min. Bacterial morphology was normal before treatment while cell destruction was observed after treatment with NB-O_3_. Scale bar, 1µm.

### Effect of NB-O_3_ treatment on total bacterial counts in fish-cultured water

In this trial, the bacterial load was compared before and after treatment. Before treatment, the total bacterial concentration in the fish-cultured water was 6.93 × 10^5^ ± 7.81 × 10^5^ CFU/mL (Fig. 6). After exposure to NB-O_3_ for 10 min, 42.94% of the bacteria was inactivated. When the same protocol was repeated, 84.94% and 99.27% bacteria were reduced in these treatments (Fig. 6).

**Figure 6:**
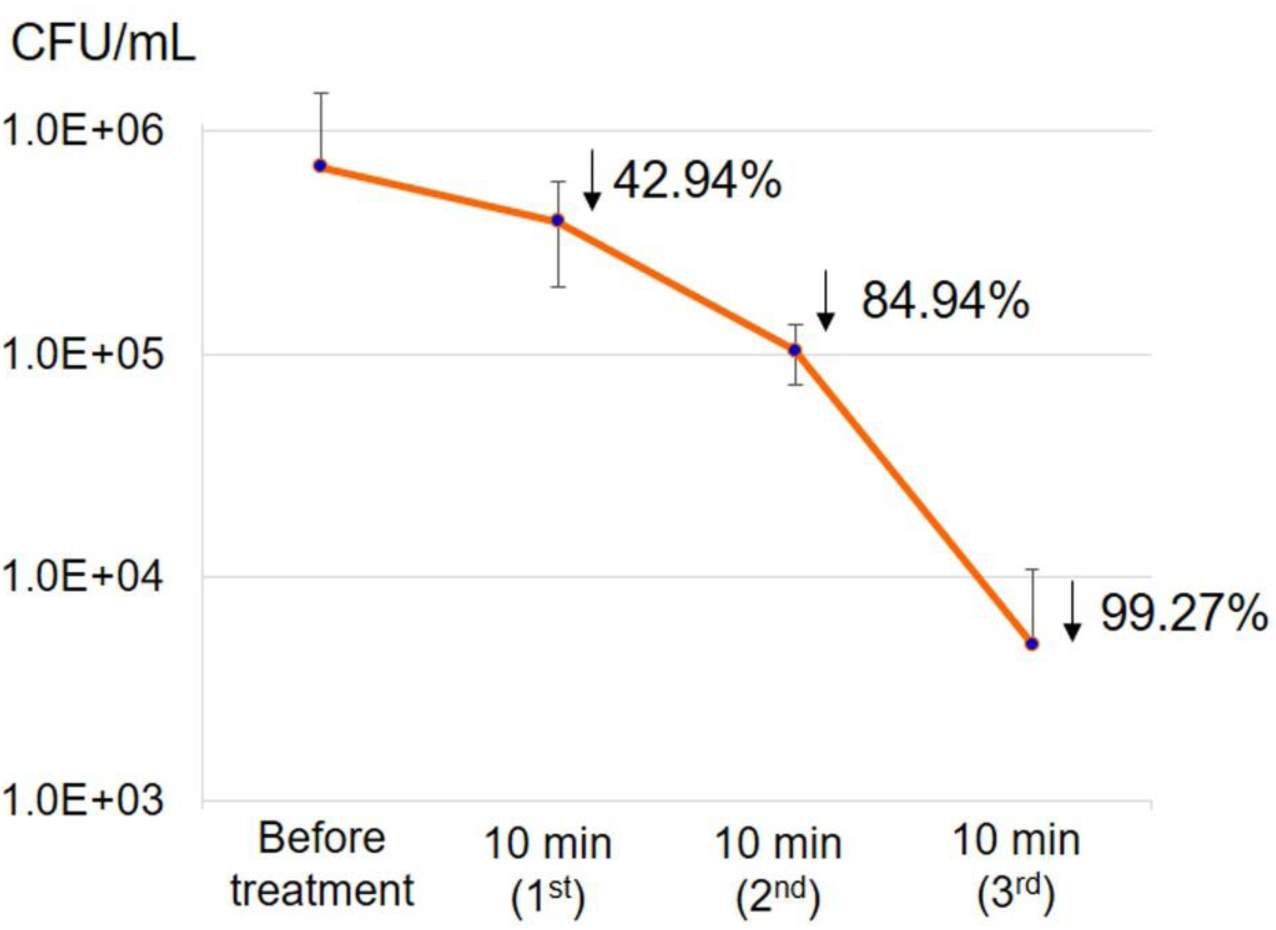
Total bacterial counts from fish-cultured water upon exposure to NB-O_3_ 10 min three times continuously. Arrows indicated % reduction of bacterial loads compared to the starting bacterial concentration. Bars represent standard deviation from 3 replicates.

During the experiment, DO increased sharply from very low at the beginning 0.6 ± 0.1 mg/L to 27.7 ± 0.6 mg/L after the first 10 min treatment. The DO was 30.8 ± 7.7 mg/L after the second 10 min treatment, and 28.7 ± 7.6 mg/L after the third treatment. Water temperature was increased slightly from 26.7 ± 0.3 to 28.3 ± 0.4, 29.8 ± 0.3 and 31.2 ± 0.2 ^°^C after the 1^st^, 2^nd^ and 3^rd^ treatment, respectively. In contrast, pH and ORP were stable during the experiment (7.5-7.6 for pH, 210-250 mV for ORP).

### Effect of NB-O_3_ on fish health and gill morphology

No Fish died during the NB-O_3_ treatments or up to 48 h post treatment when we stopped the experiment. However, abnormal signs were observed in the gills in all fish examined after receiving the second and third treatments. These signs included reddening at the base of the fins, erratic swimming, and the attachment of bubbles to the body surface. These bubbles disappeared after several minutes of fish movement.

The wet-mount examination of the gills revealed no significant difference between control and treatments at any of the treatment times (Fig. 7A-D). There were no gross clinical signs of gas bubble disease. H&E stained sections of the gills showed the normal structure of the gills in the first treatment group (Fig. 7F) compared to the control group (Fig. 7E). However, abnormal changes were observed in the fish exposed to the second treatment. Aggregates of basal cells at the base of the secondary lamellae were apparent with increasing severity corresponding to the dose of ozone exposure (Fig. 7G, arrows). Gills in the third experiment had some loss of the secondary lamella (Fig. 7H, arrows) and infiltration of red blood cells (blood congestion) (Fig. 7H).

**Figure 7:**
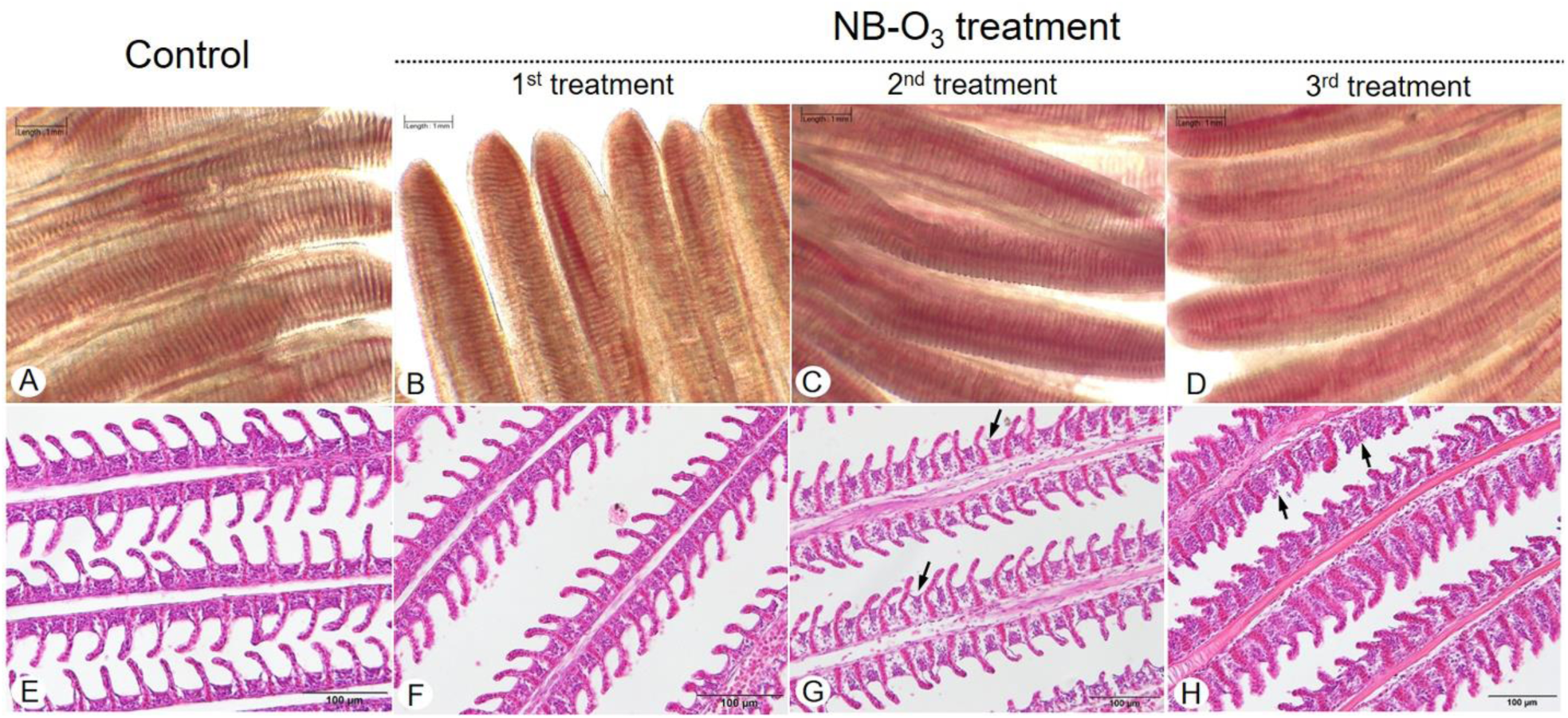
Photomicrographs of wet-mount (A-D) and H&E stained sections (E-H) of the gills of tilapia from control and NB-O_3_ treatment. No significant difference in gill morphology by wet-mount between control (A) and treatment (B-D) groups. H&E staining revealed the normal structure of the gill filaments in both control (E) and the first treatment with NB-O_3_ (F). Slight damage and shrunken of the basal lamellae (arrows) were observed in the fish received second exposure (G) and increasing damage of the gill filaments, loss of some secondary lamella (arrows) and severe blood congestion in the secondary lamellae were observed in the fish received the third exposure (H).

During the treatment, water parameter (T^°^, DO and pH) fluctuations were similar (Table 2) to the experiment with clean water spiked with *S. agalactiae* or *A. veronii* and NB-03 with the exception that tanks exposed to ozone had ORP levels of 860-885 mV after each10 min treatment.

**Table 2:** Water parameter fluctuation in fish tanks with and without and NB-O_3_ treatment.

| Parameter | Measurement time | Control | NB-O<sub>3</sub> treatment |
| --- | --- | --- | --- |
| T<sup>0</sup> | Before treatment | 28.7 ± 0.1 | 28.8 ± 0.0 |
|  | 10 min (1<sup>st</sup>) | ND | 29.6 ± 0.5 |
|  | 10 min (2<sup>nd</sup>) | ND | 30.7 ± 0.4 |
|  | 10 min (3<sup>rd</sup>) | 26.7 ± 0.1 | 31.6 ± 0.3 |
| DO | Before treatment | 4.9 ± 0.1 | 4.6 ± 0.1 |
|  | 10 min (1<sup>st</sup>) | ND | 28.2 ± 0.1 |
|  | 10 min (2<sup>nd</sup>) | ND | 28.5 ± 0.6 |
|  | 10 min (3<sup>rd</sup>) | 5.1 ± 0.0 | 26.9 ± 0.2 |
| pH | Before treatment | 8.0 ± 0.0 | 8.0 ± 0.0 |
|  | 10 min (1<sup>st</sup>) | ND | 7.6 ± 0.1 |
|  | 10 min (2<sup>nd</sup>) | ND | 7.6 ± 0.1 |
|  | 10 min (3<sup>rd</sup>) | 7.15 ± 0.2 | 7.3 ± 0.0 |
| ORP* | Before treatment | 314 ± 13 | 337 ± 6 |
|  | 10 min (1<sup>st</sup>) | ND | 860 ± 42 |
|  | 10 min (2<sup>nd</sup>) | ND | 875 ± 18 |
|  | 10 min (3<sup>rd</sup>) | 313 ± 12 | 885 ± 15 |

T<sup>o</sup>, temperature in degree Celsius; DO, dissolved oxygen; ORP, oxidation reduction potential; ND, not done. Values are expressed as mean ± SD from 3 replicates. \*ORP dropped to normal (~330 mV) after 15 min of every treatment time.

## Discussion

Application of ozone gas using nanobubble technology is relatively new to aquaculture. A previous study reported the sterilization efficacy of NB-O_3_ against pathogenic *V. parahaemolyticus*, a Gram negative bacteria causing disease in marine shrimp (Imaizumi et al., 2018). In this study, we first revealed that NB-O_3_ has disinfection property against two common bacterial pathogens of freshwater farmed tilapia, *S. agalactiae* and *A. veronii*.

The disinfection effectiveness of NB-O_3_ likely depended on the organic load in the water. In clean de-chlorinated tap water spiked with a known concentration of either *S. agalactiae* or *A. veronii*, a single treatment (10 min) with NB-03 could successfully reduce more than 96% of the bacteria. However, the same protocol applied to water that was taken from a tilapia-cultured tank, resulted in a reduction in the disinfection potential by roughly half. Ozone is known as a strong oxidizing agent (Powell et al., 2016; Summerfelt, 2003); thus, it was possible that organic matter (e.g. feces, mucus, etc.) in the dirty tank water competed for the oxidation potential of the NB-03 thus slowing down the speed of disinfection. This finding suggests that increased treatment time or increasing the frequency of treatments, as was evaluated in this study, may be required for water with abundant organic matter.

Interestingly, we also noticed that when bacteria (organic matter) was were added to water, oxidation reaction potential (ORP) value did not increase as seen in the treatment with clean water that did not have bacteria. Similarly, ORP did not increase during treatment with the fish-cultured water (rich of organic matter). This indicated that the measurement of ORP as an indicator of O_3_ level administered by the nanobubbler is not reliable in the presence of organic matter. It was probably due to the rapid oxidation and degradation of O_3_ molecules when contacting organic matters. Therefore, to accurately measure ORP in NB-O_3_ water, clean water without organic matters is required. In clean water, ORP dropped relatively quick and returned to normal after we ceased to introduce NB-O_3_ (Fig. 2), indicating that O_3_ molecules might be unstable even in the form of nanobubbles. This is consistent with the high levels of DO maintained after treatment (Fig. 2), most likely derived from the degradation of O_3_ into O_2_ molecules (Batakliev et al., 2014). If this is the case, the treatment of NB-O_3_ in aquaculture farms could have dual benefits: disinfection of bacteria and improvement of DO.

In this study, extreme treatment conditions (repeating treatments 3 times at 15 minute intervals) was designed to evaluate the acute effect of NB-O_3_ on the fish. Although multiple NB-O_3_ treatments were not harmful to fish life, increased exposure caused damage to the fish gills. In fact, a single treatment with 10-min NB-O_3_ is enough to effectively reduce bacterial loads in water, and it was safe for fish. If more than one 10-minute treatment of NB-03 was used there was some evidence of irritation to the gills. In reality, if this technology is applied in fish ponds, chances of contact between fish and NB-O_3_ will inevitably be low. However, given the evidence of gill damage after 3 consecutive treatments more in-depth investigations are required prior to scaling up NB-O_3_ technology for commercial applications.

One of the limitations of this study was the limited sample size with the experiments. Our tank numbers were limited by the number of nanobubble generators we had. Also we could not include a normal ozone air-stone treatment group due to the personnel safety issue in our laboratory. However, when we consider all the experiments together there is strong evidence to suggest that NB-O_3_ technology is not only a promising disinfection method but also enriches dissolved oxygen in freshwater aquaculture and in low dose it is not harmful to the fish. As a disease prevention tool, NB-O_3_ treatment might be a novel approach to controlling overgrowth of pathogenic bacteria in water, thus reducing the risk of bacterial diseases. This nonchemical disinfection technology may be a promising alternatives to antibiotics as a means of reducing antibiotic use in aquaculture, and possibly inadvertently reducing the risk of AMR. We are currently investigating the effect of NB-O_3_ on fish immunity and stress response, microbiome, plankton profiles and growth performance.

## Acknowledgements

This work was carried out with financial support from UK government – Department of Health and Social Care (DHSC), Global AMR Innovation fund (GAMRIF) and the International Development Research Center (IDRC), Ottawa, Canada.

## Disclaimers

The views expressed herein do not necessarily represent those of IDRC or its Board of Governors.

## Credit Author Statement

HTD, SSH: Conceptualization, Methodology. HTD, SS, SSH: Data curation, Writing-Original draft preparation. CJ, NK, NP, PS: Visualization, Investigation. HTD, WP, AT: Supervision, Validation. HTD, SS, SSH, WP, AT: Writing-Reviewing and Editing. HTD, SSH: Funding acquisition.

